# Volitional control of parieto-occipital alpha lateralization via neurofeedback does not influence auditory spatial attention

**DOI:** 10.64898/2026.04.22.720048

**Authors:** Felix Stockar, Tomas Ros, Basil Preisig

## Abstract

Auditory spatial attention, the ability to focus selectively on specific sounds while ignoring others, is crucial for everyday listening. Lateralized alpha oscillations over the parieto-occipital cortex have been proposed to act as a crossmodal attentional filter mechanism that allows to gate sensory processing by actively suppressing unattended locations. While prior evidence supports the functional role of this mechanism in visual attention, its role in auditory spatial attention remains debated. In this pre-registered EEG-neurofeedback study, participants were trained for 30 minutes to lateralize alpha power toward the left or right parieto-occipital cortex, while auditory probes from different spatial directions (-90°, -45°, 45°, and 90°) assessed whether online changes in alpha lateralization influenced auditory processing. Resting-state and task-based alpha lateralization, as well as gaze behavior were measured before and after training. Participants successfully modulated alpha lateralization in the trained direction during neurofeedback. However, shifting alpha lateralization did not cause changes in online auditory processing, as evidenced by the absence of asymmetric changes in auditory evoked responses to lateralized probe tones. Likewise, neurofeedback did not cause persistent changes in alpha lateralization during resting-state and task-based recordings after neurofeedback. Notably, neurofeedback training affected gaze behavior. Shifting alpha lateralization toward the left hemisphere during neurofeedback abolished a pre-existing rightward gaze bias in training responders, pointing to a dissociation between oculomotor and auditory attentional systems. These findings challenge the notion that parieto-occipital alpha lateralization serves as a universal crossmodal spatial gate, and raise important questions about the functional specificity of alpha-based neurofeedback interventions.

## 1 Introduction

Humans and other animals can focus on one sound among many and selectively follow it over time (Shamma et al., 2011). In everyday life, this ability is crucial for understanding speech in noisy environments, such as crowded restaurants or busy streets (Haykin & Chen, 2005). However, despite the brain’s ability to prioritize relevant auditory input, many individuals—especially those with hearing impairments—struggle to do so in complex acoustic settings (Bergman, 1971; Blumenfeld et al., 1969; Vielsmeier et al., 2016). Current hearing aids cannot selectively enhance the attended sound stream, highlighting the need for a deeper understanding of the neural mechanisms that support auditory attention (Dau & de Cheveigné, 2017).

One promising approach to investigating these mechanisms is to examine attention-related changes in neural oscillations, particularly in the alpha frequency band (8–12 Hz), which has been linked to the top-down modulation of sensory processing. Brain regions engaged in task-relevant processing typically show reduced alpha power, whereas regions associated with task-irrelevant or potentially distracting processes exhibit relatively increased alpha power (Kelly et al., 2006; Klimesch, 1997; Klimesch et al., 1999; Peylo et al., 2021; Preisig, 2024; Thut et al., 2006). This observation has led to the proposal that alpha activity serves as a gating mechanism, inhibiting task-irrelevant regions to direct information flow toward task-relevant brain regions (Jensen & Mazaheri, 2010). Notably, this gating mechanism has been proposed to operate cross-modally, with parieto-occipital alpha oscillations coordinating attentional selection across sensory modalities rather than operating exclusively within a single sensory domain (Frey et al., 2015).

In the context of spatial attention, tasks requiring spatial shifts in attention have been shown to induce hemispheric lateralization of alpha activity over occipital, parietal but also the respective sensory cortices (Haegens et al., 2011; Kerlin et al., 2010; Thut et al., 2011; Wöstmann et al., 2016). This alpha lateralization is typically characterized by an increase in oscillatory alpha power in the hemisphere ipsilateral to the attended stimulus and a decrease in the contralateral hemisphere. In the visual domain, there is evidence that alpha-band lateralization contributes functionally to the spatial deployment of attention in humans (Bagherzadeh et al., 2020; Romei et al., 2010; Thut et al., 2011). For instance, Romei et al. (2010) demonstrated that inducing alpha oscillations via repetitive transcranial magnetic stimulation (rTMS) over parieto-occipital cortex at 10 Hz selectively impaired visual detection in the hemifield opposite to the stimulated hemisphere, whereas stimulation at beta (20 Hz) or theta (5 Hz) frequencies did not. Moreover, concurrent EEG recordings have shown that rTMS applied at a participant’s individual alpha frequency over the parietal regions induces and enhances alpha activity (Thut et al., 2011). However, the functional relevance of parieto-occipital alpha activity in the auditory domain remains debated (Dahl et al., 2019; Getzmann et al., 2020; Tune et al., 2021). While some studies report no significant relationship between parieto-occipital alpha lateralization and auditory attention (Klatt et al., 2020; Tune et al., 2021), others found a correlation (Dahl et al., 2019).

The aim of this study was to test whether parieto-occipital alpha activity serves as a cross-modal attentional mechanism that contributes functionally to auditory spatial attention. Previous studies applying non-invasive electric brain stimulation with alternating currents in the alpha frequency band to parietal (Deng et al., 2019) and temporoparietal regions (Wöstmann et al., 2018) suggest a causal link between auditory attention and alpha-band activity. However, while these studies provide behavioral evidence, they cannot directly relate changes in auditory spatial attention to underlying neural alpha lateralization, as transcranial alternating current stimulation (tACS) introduces substantial artifacts into the EEG signal (Noury et al., 2016; Noury & Siegel, 2018). To overcome this limitation, the present study used neurofeedback to experimentally manipulate alpha lateralization. This approach allowed us to assess its effects on auditory sensory processing and selective attention while simultaneously measuring changes in neural alpha lateralization.

Neurofeedback is a technique that enables individuals to regulate their own brain activity using real-time sensory feedback (for reviews, see (Ros et al., 2014; Sitaram et al., 2017; Thibault et al., 2016)) In humans, neurofeedback can be based on non-invasive measurements of neural activity obtained through EEG, MEG, or fMRI. For example, Ros et al. (2010) found that the endogenous suppression of alpha oscillations via neurofeedback leads to changes in motor-evoked potentials elicited by TMS. Previous research has demonstrated that neurofeedback can be used to train participants to modulate their neural alpha lateralization, which caused a spatial bias in visual processing and visual spatial attention (Bagherzadeh et al., 2020; Okazaki et al., 2015). Thus, neurofeedback could be a promising approach to study the functional role of alpha oscillations in auditory attention.

In the present study, we used EEG-based neurofeedback to test whether changes in alpha lateralization have immediate (online) effects and longer-lasting (offline) effects on auditory spatial attention. In a multi-session cross-over design, participants were trained to increase alpha power over the left relative to the right parieto-occipital cortex (left neurofeedback training, LNT) and vice versa (right neurofeedback training, RNT). Neurofeedback was provided in the form of a computer game: when participants modulated their alpha power in accordance with the training condition (LNT or RNT), their spaceship accelerated; modulation in the opposite direction reduced the spaceship’s speed. The online effects of neurofeedback on auditory spatial attention were tested by presenting auditory probe sounds from various horizontal locations during the neurofeedback sessions. To assess offline effects, we measured alpha lateralization before and after training using electroencephalography (EEG) recording at rest and during a spatial pitch discrimination task (Wöstmann et al., 2019). To further assess cross-modal effects of neurofeedback on visual attention, we additionally evaluated the influence of neurofeedback training on participants’ gaze direction throughout the entire experiment. This was motivated by evidence that attentional shifts, including covert ones, are accompanied by shifts in eye gaze in the form of microsaccades (Liu et al., 2022a, 2023), and that auditory attention can influence gaze direction and modulate retinotopic activation in a manner similar to visual attention.

## 2 Methods

The design and analysis plan were preregistered on the Open Science Framework (https://doi.org/10.17605/OSF.IO/D7WGM), and the experiment was conducted in accordance with this preregistration. Any minor adjustments or exploratory analyses are explicitly listed and justified.

## 3 Participants

Thirty-five volunteers (*M* = 21.6 years, *SD* = 2.3, age range 18–28 years, 5 males) participated in the study. All participants were Swiss German speakers, right-handed, and had normal hearing and vision. They reported no history of neurological or psychiatric disorders. Further, all participants were monolingually raised, had no history of dyslexia, and reported no current use of medications with cognitive side effects or psychoactive substances. Hearing ability was assessed with pure tone audiometry (PTA) to determine hearing levels from 0.25 to 8 kHz. A visual *n*-back task was administered to assess the working memory capacity of participants. Additionally, handedness was assessed using the Edinburgh Handedness Inventory (EHI, Oldfield, 1971). Participants were recruited via the institute’s mailing list. They were compensated with 25 CHF per hour or course credits for their participation. All participants gave written informed consent prior to the experiment. The study was conducted in adherence to the Declaration of Helsinki and received approval from the local ethics committee (Kantonale Ethikkommission Zürich, BASEC2022-01164)

### 3.1 Procedure

The study employed a within-subjects crossover design comprising the two experimental training conditions, left neurofeedback training (LNT) and right neurofeedback training (RNT). The experiment was divided into two sessions on separate days. The average time interval between the first and second session was *M* = 10.92 days (range: 1–27). The order of the training conditions was counterbalanced across participants: those with an odd participant ID were assigned to start with LNT and those with even ID to start with RNT. The overall experimental procedure was consistent across both sessions. Following the screening, which included PTA, the visual n-back task and the Edinburgh Handedness Inventory, we evaluated alpha power lateralization and auditory attention during a spatial pitch discrimination task (see dedicated section below). The task had a duration of approximately 45 minutes. Afterwards, a two-minute resting-state EEG recording was conducted with participants’ eyes open followed by the NF training. To test the online effect of NF on auditory spatial attention, auditory probes were presented from four loudspeakers (AURATONE 5C Super Sound Cube) positioned at - 90°, -45°, +45°, and +90° on the horizontal axis. By varying the spatial locations of the probes, we aimed to assess whether training-related changes in alpha power lateralization influenced the magnitude of event-related potentials (ERPs) based on the tone’s horizontal position. NF was subdivided into four blocks of equal length and lasted around 30 minutes. After NF training, another two-minute resting-state EEG was recorded, followed by a post measurement of auditory attention and alpha lateralization using the same pitch discrimination task (see Figure 1). The second session concluded with a debriefing. The experiment was conducted in a sound-attenuated cabin. Throughout all phases of the experiment (task, resting-state EEG, and NF), eye movements were recorded. Each session had a duration of 3 to 3,5 hours.

**Figure 1.**
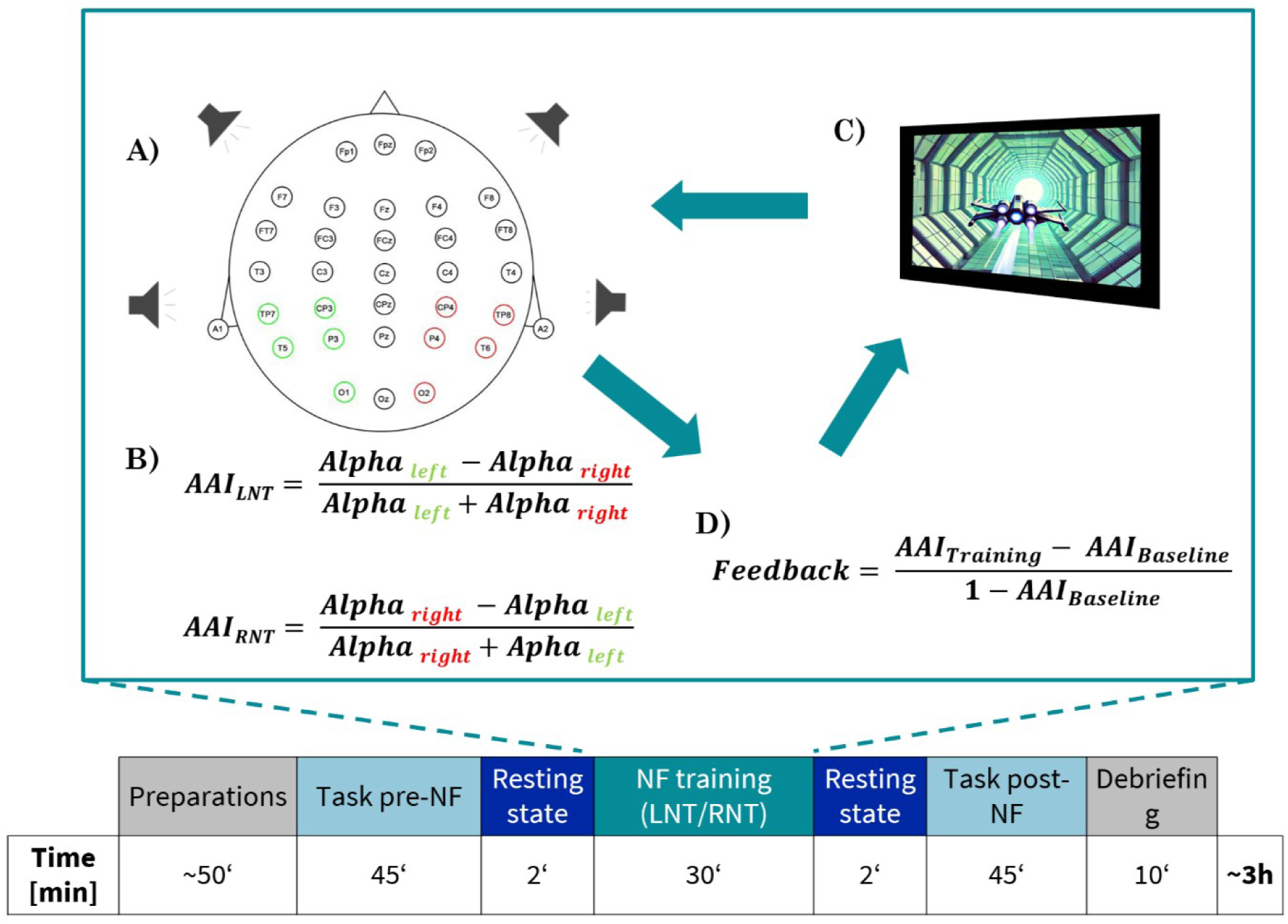
Overview of the neurofeedback (NF) training procedure and computation of the feedback signal. (A) Analysis of the EEG signal: Measurement of neural activity from bilateral parieto-occipital EEG channels. Left-hemispheric electrodes (green) and right-hemispheric electrodes (red) were used to extract alpha power. (B) Computation of the Alpha Asymmetry Index (AAI), which reflects lateralized alpha power and was calculated separately for each training condition (LNT; RNT). For left NF training (LNT), the formula emphasizes increased alpha in the left hemisphere; for right NF training (RNT), the formula is inverted. (C) Real-time visual feedback interface: participants controlled the speed of a virtual spaceship based on moment-to-moment changes in lateralized alpha activity. (D) Feedback values were computed relative to a baseline, using the normalized difference between AAI during training and resting state. The lower panel summarizes the timeline and duration of each experimental phase.

### 3.2 EEG recording

EEG data was recorded with a NeuroAmp II device (BEE Medic) equipped with 33 passive Ag/AgCL electrodes at a sampling rate of 500Hz. Channels A1 and A2 (left and right ear tips) served as reference electrodes, with all electrode impedances maintained below 20 kOhm.

### 3.3 Neurofeedback Training

NF was presented as a computer game in which participants controlled the speed of a spaceship by modulating their alpha power according to the training condition (LNT; RNT). Increasing alpha power in the target direction accelerated the spaceship, while modulation in the opposite direction slowed it down. The training protocol was implemented using BEE Lab Software (BEE Medic, 2023), Version 1.0.2.54) and visualized with “Inner Tube” (SomaticVision, 2012). The Alpha Asymmetry Index (AAI) was computed as the difference in alpha power between ipsilateral (IL) and contralateral (CL) EEG channels, relative to the training direction, normalized by their sum (see figure 1C, AAI range -1 to +1).

In the LNT condition, the ipsilateral channel included the left-hemispheric electrode positions TP7, CP3, T5, P3, and O1, the contralateral channel the right-hemispheric electrode positions TP8, CP4, T6, P4, and O2, and vice versa for RNT condition (see Figure 1A)

During NF, the AAI was calculated as follows. First, the EEG data was re-referenced to the average of all electrodes. Each EEG data segment was then bandpass filtered (2nd order Butterworth filter) within the alpha frequency band (8-12 Hz). Subsequently, the amplitude of the signal envelope was determined for both channel groups (IL; CL) using a sliding 200 ms time window, and the corresponding AAI was computed. Feedback, represented by the spaceship’s speed (see figure 1D, feedback range 0 to +1 if AAI_Training_>AAI_Baseline_), was derived by normalizing the AAI during training (AAI_Training_) with the baseline AAI (AAI_Baseline_), which reflected each participant’s inherent AAI measured during the resting-state EEG before NF.

If AAI_Training_ exceeded AAI_Baseline_, the speed of the spaceship’s speed was adjusted linearly based on the resulting value, ranging from 0 (minimum speed) to 1 (maximum speed). Conversely, if AAI_Training_ was lower than AAI_Baseline_, resulting in a negative value, the spaceship stopped. Participants were instructed to maintain their focus on the spaceship, which was presented in the center of the screen.

To test the online effect of NF on spatial auditory attention, auditory probes were presented from four loudspeakers (AURATONE 5C-super-sound-cube) positioned at -90°,-45°,+45°, and +90° degrees, aligned with the participants’ ears and positioned at 70 cm relative to the participant’s vertex. The speakers were powered by three amplifiers (Crown XL5 402), and an RME Fireface UFX II soundcard was used for the multi-channel setup. The auditory probes were presented with a randomly jittered inter-stimulus interval ranging from 3 to 4.5 seconds. A total of 400 probes were presented during NF training, evenly distributed across the four horizontal directions.

The training was divided into four blocks, each containing 100 probes (25 per direction, randomly distributed within each block) and lasting approximately 7 minutes, depending on the inter-stimulus interval. A one-minute break was provided between blocks.

### 3.4 Pitch discrimination task

To assess potential changes in alpha lateralization induced by NF training, we used a spatial pitch discrimination task developed by Wöstmann et al. (2019). This task allows to evaluate alpha lateralization based on both lateral target selection and lateral distractor suppression.

The participants listened to two concurrently presented tone sequences, presented either from the left side and the front (-90°, 0° azimuth relative to ear-nose-ear line) or from the front and the right side (0°, 90° azimuth). Each tone sequence comprised two complex tones with different pitches (low- versus high). The tones had a duration of 0.5 seconds. At the beginning of each trial, an auditory spatial cue was presented on one of the loudspeakers (front, left, or right) to indicate the position of the target tone sequence. The cue was followed by a jittered cue-to-target interval averaging 1.8 seconds (range: 1.47–2.48 s). The tones were then presented with a fixed inter-stimulus interval of 50ms. To maintain task difficulty, pitch discrimination accuracy was adaptively tracked using a 1-up-2-down staircase procedure. In this procedure, the high-pitch tone was adjusted by 0.1 semitones to maintain performance at approximately 71% accuracy within each block. Each session consisted of 8 task blocks: four before NF training (blocks 1-4) and four after (blocks 5-8).

Participants had to report whether the pitch of the attended tone sequence was increasing (low tone- high tone) or decreasing (high tone – low tone) and their confidence in this decision. Responses were recorded using a 4-button response box (MilliKey, SR-5 r2) with options for rising tone high confidence, rising tone low confidence, falling tone low confidence, and falling tone high confidence. The order of blocks was counter-balanced across participants and alternated between left and right. Within each block, participants completed 64 trials.

In summary, the task manipulated two independent variables: the location of the lateral stimulus (left side vs. right side) and the position of the target stimulus (front vs. side), resulting in four experimental conditions:

- Attend to front, ignore left.
- Attend to front, ignore right.
- Ignore front, attend to left.
- Ignore front, attend to right.

The task was programmed by Wöstmann et al. (2019) using Psychtoolbox (Brainard, 1997) in MATLAB.

### 3.5 EEG Preprocessing

The raw eye-tracking and EEG data were synchronized using the EEGLab toolbox (Delorme & Makeig, 2004). Thereby, eye-tracking data were automatically downsampled from 1000 Hz to 500 Hz to match the sampling frequency of EEG data. The synchronized dataset consisted of 31 EEG channels and 8 eye tracking channels. Further preprocessing was conducted using the fieldtrip toolbox (Oostenveld et al., 2011). EEG data recorded during task, resting-state and NF were preprocessed as follows: continuous EEG data was segmented into 1-second epochs. Data cleaning involved visually identifying and excluding noisy epochs and channels. Bad channels were interpolated, and epochs with voltage deviations exceeding ±150 μV at any of the electrodes were rejected. Next, EEG data was bandpass filtered between 2 and 40 Hz and re-referenced to the average of all electrodes. Ocular artifacts were removed using independent component analysis (ICA). Blink- and horizontal eye movement-related components were identified through visual inspection, with one to two components removed per participant. After ICA, preprocessed EEG data were downsampled to 250Hz and into a single continuous dataset.

### 3.6 EEG offline Analyses

#### 3.6.1 Neurofeedback and resting-state spectral analysis

To assess whether participants modulated their alpha lateralization during neurofeedback, preprocessed EEG data from training and pre- and post-resting-state sessions were analyzed in 1-second segments. To minimize contamination of the neurofeedback data by auditory ERPs, the first 500 ms following each auditory probe stimulus were excluded from analysis before segmentation. Power spectral density (PSD) was estimated using a multi-taper fast Fourier transformation. The analysis was conducted separately for left and right electrode clusters (see Figure 1A). To improve spectral resolution, the data were zero-padded to the next power of two, and power estimates were obtained in the 2–30 Hz frequency range. Afterwards, the AAI was computed for the alpha frequency range (8–12 Hz) based on the PSD from EEG channels over the left and right hemispheres. PSD values were extracted separately for left and right hemisphere electrodes, and the AAI was calculated as the normalized difference (see formula 1) between left and right hemisphere channels. This analysis was conducted separately for pre-NF resting-state EEG (pre-NF rest), NF training itself, and post-NF resting-state EEG (post-NF rest).

#### 3.6.2 Probe-related auditory evoked responses

To test the online effect of NF training on auditory spatial attention, preprocessed EEG data for each participant were segmented into event-related trials from -1 to 3s relative to probe onset. The primary time window of interest was 0–500 ms post-stimulus. First, the grand-average ERP was computed across all conditions and participants to examine the overall waveform shape. Based on the resulting ERP topography (Figure 2), a set of vertex channels (FC3, FCz, FC4, C3, Cz, C4) was selected for further analysis. For each participant and stimulus condition (stimulus direction: -90°, -45°, 45°, 90°), ERPs were calculated across the above-defined vertex channels. The average ERP across all trials was computed for each condition and participant. The ERP amplitude was quantified as the absolute difference between the maximum value in the P1 window (50–80 ms) and the minimum value in the N1 window (80–130 ms). These ERP amplitudes were calculated for each condition and participant.

**Figure 2:**
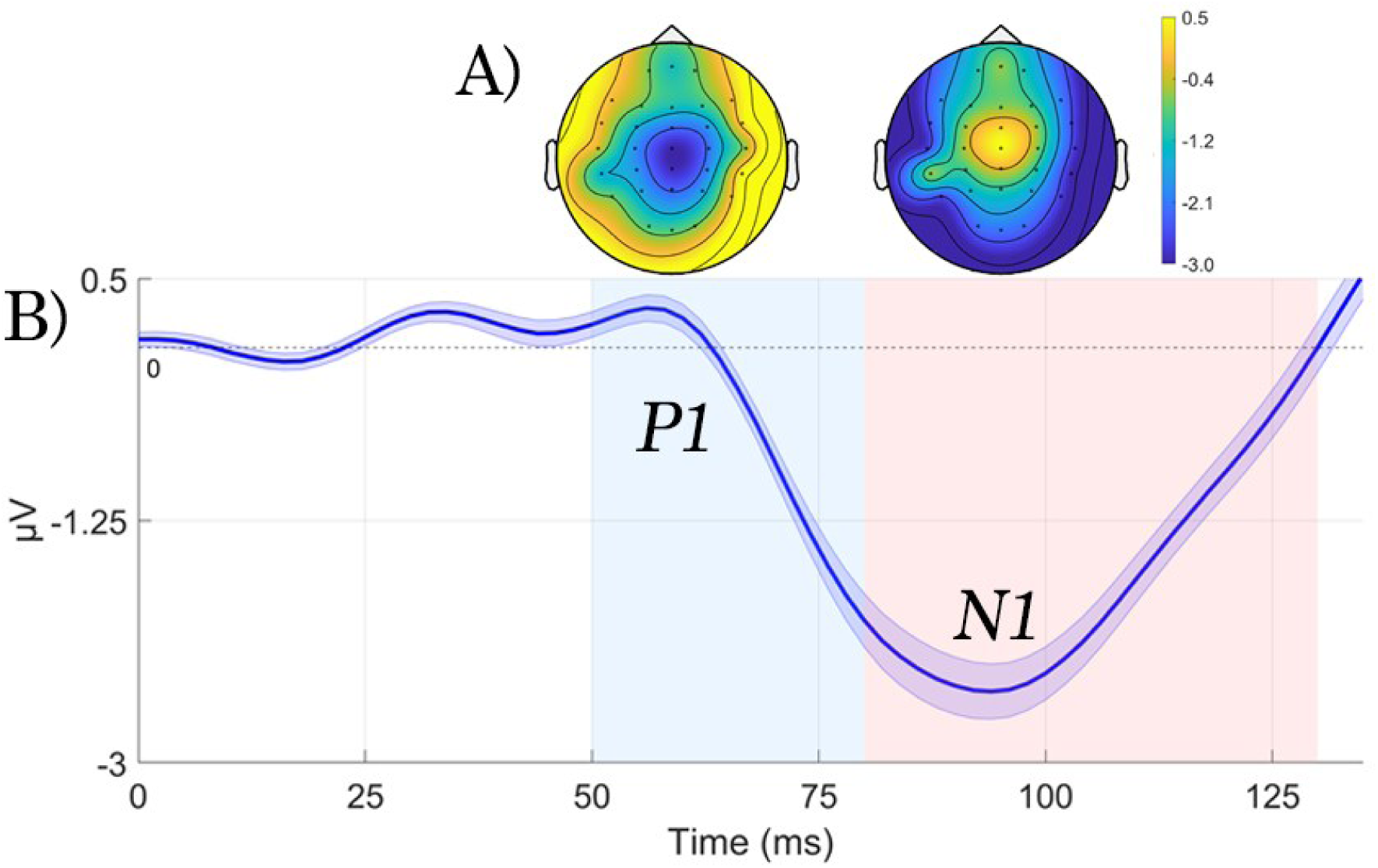
Grand-average event-related potentials (ERPs) across all conditions and participants. (A) Topographical maps for the P1 (50–80 ms) and N1 (80–130 ms) components. Color coding represents ERP amplitudes, with positive values (yellow) indicating stronger activation and negative values (blue) indicating lower activation. (B) ERP waveform averaged across participants, channels and spatial positions of the auditory probes. The shaded region represents the standard error of the mean (SEM). Light blue and red background shading indicate the predefined time windows for the P1 and N1 components.

#### 3.6.3 Task

First preprocessed EEG data were segmented into epochs from -2 to 6 seconds relative to the onset of the auditory spatial cue that indicated the position of the to be attended tone sequence. Time-frequency representations (TFRs) were computed using a multi-taper convolution method for the 2-30 Hz frequency range. TFRs were calculated separately for the four experimental conditions. and analyzed independently for pre- and post-NF task blocks. To assess the effects of lateral target selection and lateral distractor suppression, we computed how alpha power spectral density was modulated. For this purpose, the condition dependent alpha modulation index (AMI) was calculated separately for conditions involving lateral target selection (select-left; select-right) and distractor suppression (suppress-left; suppress-right). For each participant and EEG-channel, two AMIs were computed based on absolute oscillatory power, one quantifying target selection (AMI_Selection_), the other quantifying distractor suppression (AMI_Suppression_).

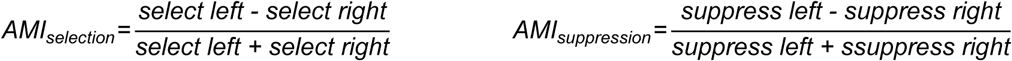

The analysis was based on two sets of left-hemispheric electrode positions (TP7, CP3, T5, P3, and O1), and right-hemispheric electrode positions (TP8, CP4, T6, P4, and O2). For each of these sets the AMI was computed based on the average power in the alpha band (8 –12 Hz) within the time interval between the onset of the spatial cue and the onset of the tone sequences (0 –1.8 s). Wöstmann et al. showed that alpha power is modulated in opposite directions during lateral target selection and lateral distractor suppression, i.e., alpha power is increasing ipsilateral to the presented targets and contralateral to the presented distractors.

The AAI was then computed as the difference in the AMI between the left and the right electrode sets:

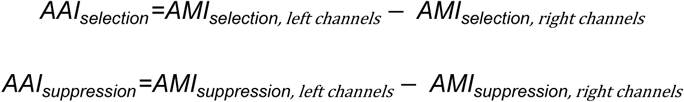

### 3.7 Eye-tracking recording

Throughout the entire experiment eye movements were recorded with an EyeLink-1000 Plus eye-tracker (SR Research). Eye movements were recorded from an eye distance of ∼50cm, using binocular pupil tracking at 1000 Hz. Participants placed their head on a height-adjustable chin rest. A standard 9-point calibration procedure (Tatler et al., 2005) was conducted before the first resting state recording and again before NF-training.

### 3.8 Eye-tracking analysis

The aim of the eye-tracking analysis was to examine whether gaze behavior mirrored NF-induced shifts in alpha lateralization, specifically by analyzing changes in gaze bias before and after NF training and comparing gaze behavior between the two NF training conditions.

Eye-tracking data were synchronized with EEG data (see EEG Preprocessing) to ensure precise alignment of eye movement events. Like the analysis of the NF EEG data, the first 500 ms of eye-tracking data following each auditory probe stimulus were excluded to remove gaze shifts triggered by auditory probes.

Blink artifacts from both eyes were identified using pupil area signals, and time segments containing blinks were excluded. The horizontal gaze position (X-axis) was extracted separately for the left and right eye and averaged. In cases where one eye had reduced data quality (e.g., due to loss of tracking) only the data from the better eye were used. Notably, eye-tracking quality differed between datasets: for neurofeedback (NF) recordings, binocular tracking was available in 98% of the data segments, while for the auditory task (TASK) recordings, this was the case in only 48% of segments, due to different task demands.

Missing values in gaze position were linearly interpolated, and the signal was detrended to remove slow drifts. Screen coordinates (pixels) were converted to degrees of visual angle.

Microsaccades were defined as small eye movements spanning 0.03° to 0.5° visual angle (Liu et al., 2022). The microsaccade rate (Hz) was computed separately for eye movements directed toward the left and right visual hemifields, allowing us to assess whether NF training led to systematic shifts in gaze bias in line with alpha lateralization changes observed in the EEG data.

### 3.9 Statistical Analyses

EEG and Eye-tracking data processing were carried out in MATLAB (R2022a) and statistical analyses were conducted in JASP (Jasp Team, 2025). Repeated measures ANOVAs were fitted to the data. In cases where the assumption of sphericity was violated, the Greenhouse-Geisser correction (Greenhouse & Geisser, 1959) was applied, provided that the repeated measure variable had more than two factor levels. To compare individual factor levels, post hoc comparisons were conducted using paired t-tests, and Holm corrections for multiple comparisons were applied (Holm, 1979). Bayes factors were calculated based on Bayesian repeated measures ANOVAs, using the mean values of the dependent variable for each group. A Bayes factor (BF_10_) greater than 10 strongly supports the alternative hypothesis, while a BF_10_ smaller than 0.1 provides strong evidence for the null hypothesis. (Jeffreys, 1961).

## 4 Results

### 4.1 Participants modulate alpha power lateralization during Neurofeedback

First, it is important to note that positive AAI values indicate greater alpha power lateralization toward the left hemisphere, while negative values indicate lateralization toward the right hemisphere. In the current sample, participants exhibited a consistent negative lateralization in the resting-state EEG before neurofeedback (*M* = -0.095 *SEM* = 0.022). To test whether neurofeedback modulated alpha power lateralization a repeated-measures ANOVA was conducted with Time (pre-training rest; training; post-training rest) and Training (LNT; RNT) as within-subject factors, and AAI as the dependent variable. Results revealed a significant Time x Training interaction (*F*(2,68) = 7.52, *p* = 0.001, *η^2^_p_* = 0.181, *BF_10_* = 56.417), indicating that alpha power lateralization was influenced by Training (LNT; RNT) over time (see Figure 3). Pairwise comparisons were conducted to examine whether neurofeedback shifted alpha lateralization in the trained direction. In both Training conditions, the alpha lateralization index changed as expected. In the LNT condition, AAI values increased during neurofeedback, indicating a shift in lateralization towards the left hemisphere (pre-training rest: *M* = -0.09, *SEM* = 0.02; training: *M* = -0.06, *SEM* = 0.02). In the RNT condition, AAI values decreased compared to the resting-state baseline, suggesting a shift in lateralization towards the right hemisphere (pre-training rest: *M* = -0.08, *SEM* = 0.01; training: *M* = -0.11, *SEM* = 0.01). However, neither of these changes relative to baseline reached statistical significance (paired *t*-tests: LNT: *t*(34) = -1.65, *p_holm_* = 0.215; RNT: *t*(34) = 1.78*, p_holm_* = 0.195).

**Figure 3.**
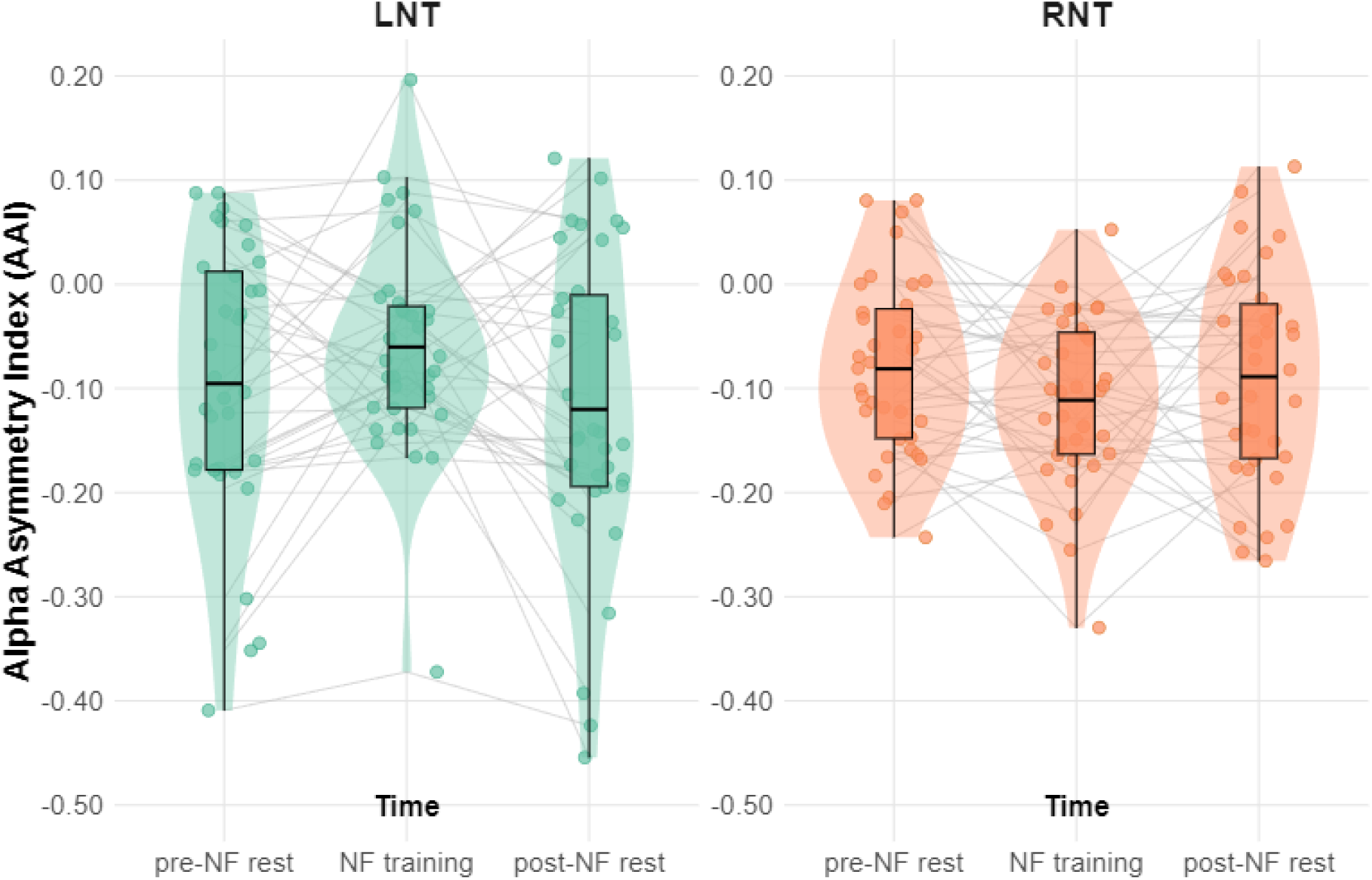
The Alpha Asymmetry Index (AAI) at three time points (pre-training rest, training, post-training rest) for the two training conditions (LNT; RNT). Positive AAI values indicate greater alpha power over the left hemisphere, while negative values indicate greater alpha power over the right hemisphere. A significant Time × Training interaction (*F*(2,68) = 6.36, *p* = 0.001, *η_2p_* = 0.181, *BF*_10_ = 56.417) shows that the direction of alpha power lateralization changed over time depending on the training condition. In both conditions, the mean AAI values shifted in the expected direction. However, pairwise comparisons were not significant for either the LNT condition (*p_holm_* = .108) or the RNT condition (*p_holm_* = .196). The middle line in each boxplot shows the median; the box represents the interquartile range (IQR, Q1–Q3), and the notches or error bars indicate the approximate 95% confidence interval (CI) of the median. The violin plot outlines show the distribution of AAI values, estimated using kernel density.

Further, we examined whether neurofeedback had lasting effects on lateralization in the post-training resting-state EEG. Our results show no persistent change in alpha lateralization. Group means in the LNT condition (pre-training rest: *M* = -0.09, *SEM* = 0.02; post-training rest: *M* = -0.11, *SEM* = 0.02) and in the RNT condition (pre-training rest: *M* = -0.08, *SEM* = 0.01; post-training rest: *M* = -0.07, *SEM* = 0.01) returned to pre-training levels (see Figure 3). In the LNT condition, 23 out of 35 participants modulated the AAI in the trained direction, compared to 21 out of 35 participants in the RNT condition. In addition, 15 participants showed modulation in both conditions, which is less than half of the sample, while 6 participants showed no modulation in either the LNT or RNT condition. Our results suggest that most participants were able to modulate alpha lateralization in one direction, but not in both. This appeared to be independent of whether they were in the first or the second session, as 24 participants modulated the AAI during the first session (15 in LNT; 9 in RNT) and 20 participants during the second session (8 in LNT; 12 in RNT).

We further examined whether online changes in alpha lateralization during neurofeedback were related to offline changes in lateralization from pre to post neurofeedback. To this end, we computed each participant’s online effect as the difference in AAI between the first and last neurofeedback block (see Figure 4A for an illustration of how AAI evolved over the course of neurofeedback), and correlated this with the pre to post neurofeedback AAI difference as a measure of the offline effect (see Figure 4B).Interestingly, the direction of this relationship differed between training conditions. In the LNT condition, the correlation was positive, suggesting that participants who showed greater leftward lateralization during neurofeedback also tended to show greater leftward offline changes, although this effect did not reach significance (*rho* = 0.19, *p* = .275). In the RNT condition, however, the correlation was negative, with a statistical trend indicating that larger rightward shifts during neurofeedback were associated with smaller offline changes in the same direction (*rho* = -0.29, *p* = .094).

**Figure 4.**
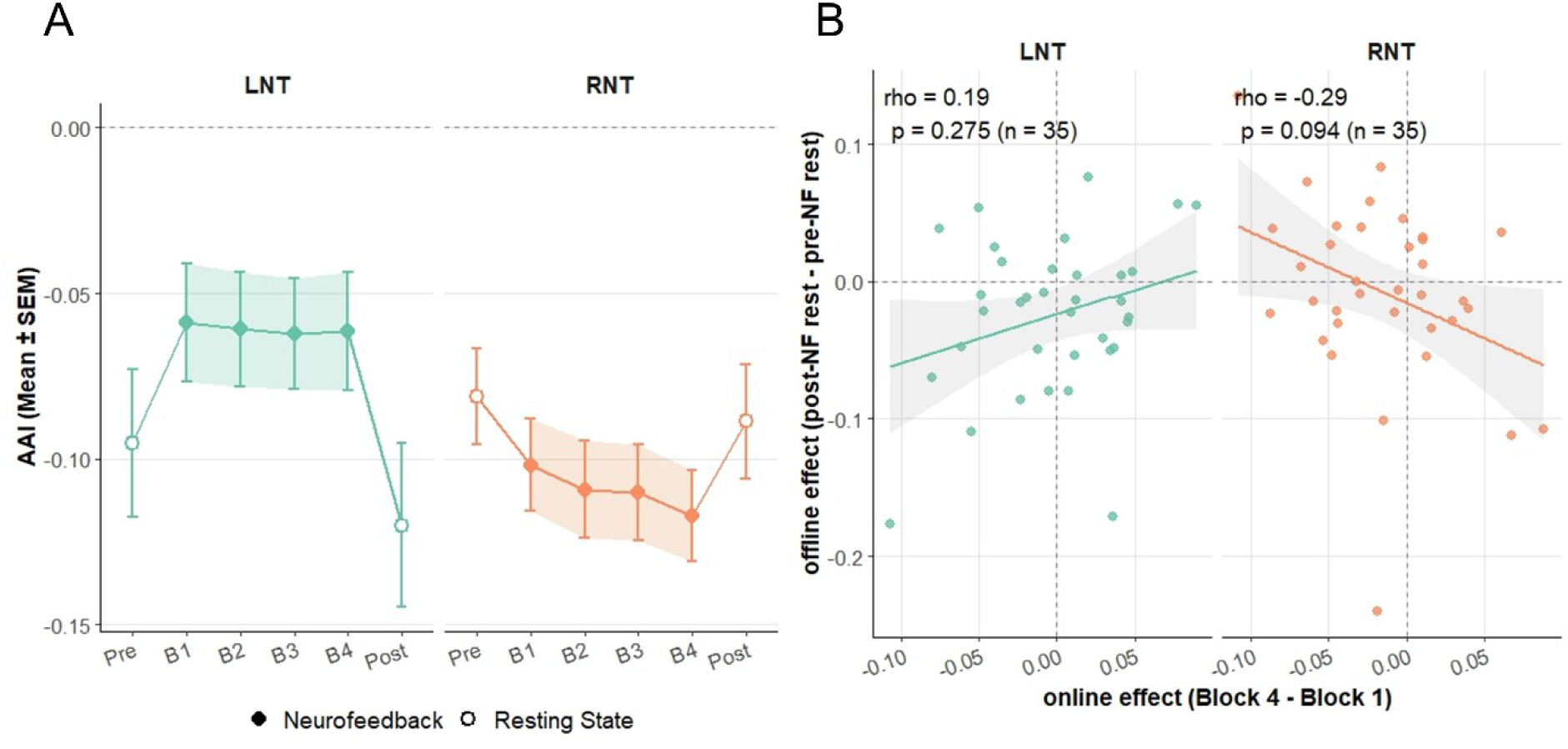
Online neurofeedback effects and their relationship to offline alpha lateralization changes. A) Evolution of the Alpha Asymmetry Index (AAI) for the two training conditions (LNT; RNT) over the experimental session. Filled circles show mean ± SEM AAI during neurofeedback blocks, empty circles show mean ± SEM AAI at rest. B) Correlation between the online and offline neurofeedback effects for both training conditions. In the LNT condition, participants who showed greater leftward lateralization during neurofeedback also tended to show greater leftward offline changes (rho = 0.19, p = .275). In the RNT condition, however, larger rightward shifts during neurofeedback were associated with smaller offline changes in the same direction (rho = -0.29, p = .094).

### 4.2 Neurofeedback does not bias online auditory processing

To test the online effect of neurofeedback on auditory spatial attention, we examined whether the training condition (LNT; RNT) influenced auditory evoked potentials elicited by probe tones presented from different spatial directions (-90°; -45°; 45°; 90°). This was tested using a repeated-measures ANOVA with within-subject factors Training (LNT; RNT) and Probe Direction (-90°; -45°; 45°; 90°) on ERP amplitude. Contrary to our hypothesis, we found neither evidence for a main effect of Training (*F*(1,33) = 0.93 *p* = 0.341, *η^2^_p_* = 0.028*, BF_10_* = 0.471), nor for a Training x Probe Direction interaction (*F*(3,99) = 0.46, *p* = 0.711, η*²_p_* = 0.014, *BF_10_* = 0.063) (Figure 5).

**Figure 5.**
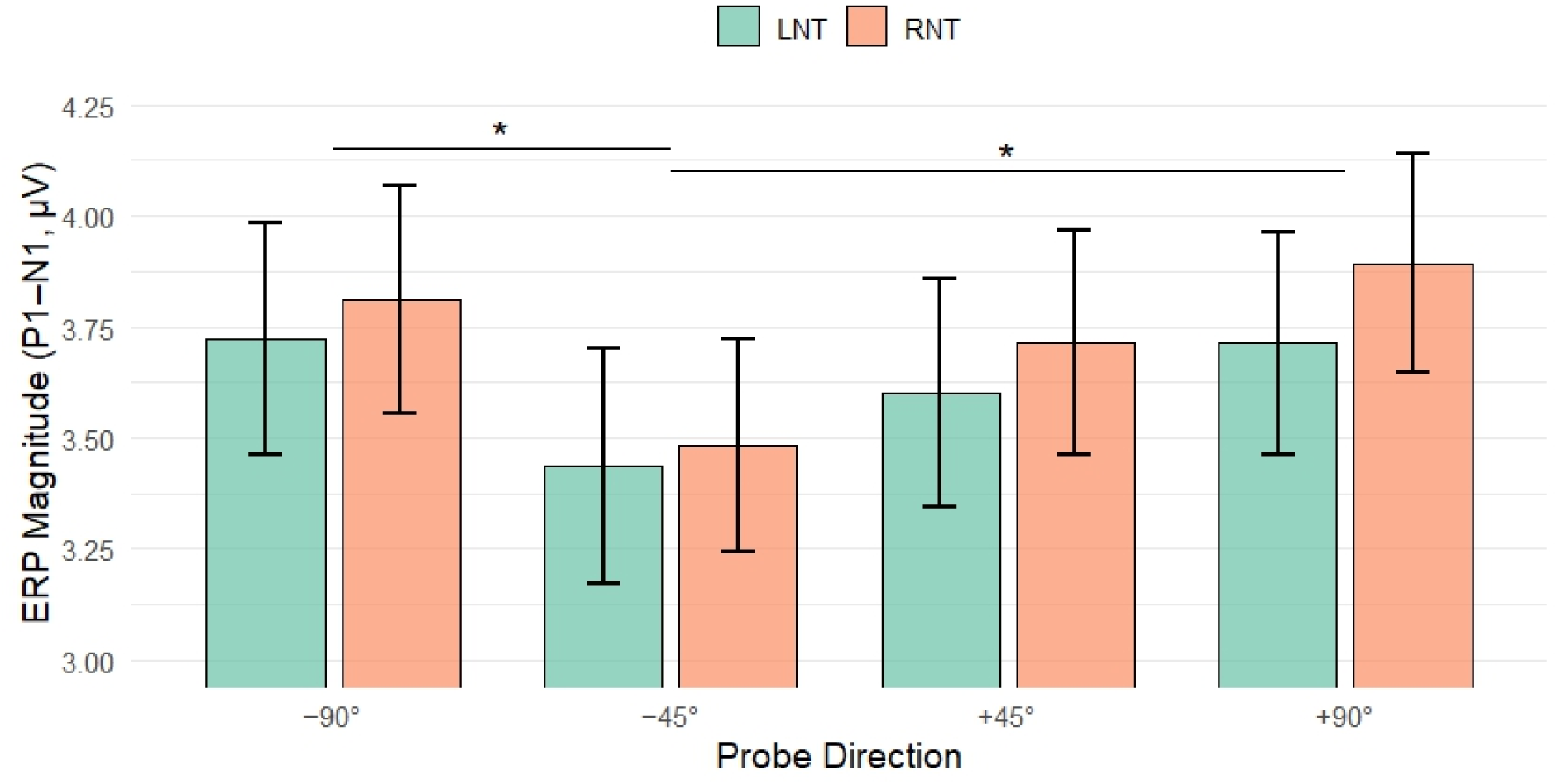
ERP magnitudes as a function of Probe Direction. ERP magnitudes (P1–N1) are shown for auditory probes presented from four horizontal directions (−90°, −45°, +45°, +90°), separately for the two training groups (LNT; RNT). A repeated-measures ANOVA revealed a significant main effect of probe direction, with reduced ERP amplitudes for probes presented at −45° compared to those presented at −90° and +90°. Horizontal lines with asterisks indicate these significant pairwise differences. Error bars represent the standard error of the mean (SEM).

Further, the analysis revealed a significant main effect of Probe Direction (*F*(3,99) = 7.505, *p* < 0.001, *η^2^_p_* = 0.185*, BF_10_* = 146.23). Post hoc analyses indicated a gradual decrease in ERP amplitude from +90° (*M* = 3.86 µV, *SEM* = 0.24) to −45° (*M* = 3.54 µV, *SEM* = 0.23; p = .003, *BF₁₀* = 117; see Figure 5). Notably, the −90° condition deviated from this pattern, showing higher amplitudes (M = 3.86 µV, SEM = 0.24) than −45° (*p* < .001, *BF₁₀* = 403577), thereby interrupting the otherwise graded trend. This spatial gradient in ERP amplitude may reflect the influence of a pre-existing rightward alpha asymmetry on auditory cortical excitability, independent of neurofeedback condition. Therefore, in a next step, we pooled ERP amplitudes across probe positions within each hemifield (left = −90°/−45°; right = +45°/+90°). A repeated-measures ANOVA with Training (LNT; RNT) and Hemifield (left; right) as within-subjects factors revealed no significant main effect of Training (F(1, 33) = 0.93, p = .341, *η^2^_p_* = .028, *BF₁₀* = 0.53), and no Training x Hemifield interaction (*F*(1, 33) = 0.63, *p* = 0.432, *η^2^_p_* = .019, *BF*₁₀ = 0.32). The main effect of Hemifield was non-significant, with inconclusive Bayesian evidence (*F*(1, 33) = 2.79, *p* = 0.104, *η^2^_p_* = .078, *BF₁₀* = 1.35).

As not all participants were able to modulate their alpha lateralization through neurofeedback, we additionally conducted exploratory analyses restricted to responders. To examine whether probe tones from different spatial directions (−90°, −45°, 45°, 90°) modulate ERP amplitudes in neurofeedback responders, we conducted exploratory repeated-measures ANOVAs separately for the LNT and RNT responder groups, with Probe Direction (−90°, −45°, 45°, 90°) as a within-subject factor. Given that rightward alpha lateralization was more pronounced in the RNT condition, a stronger effect of Probe Direction was expected in RNT responders. For LNT responders (n = 23), the main effect of Probe Direction was significant (*F*(3, 63) = 3.87, *p* = 0.013, *η^2^* = .150, *BF₁₀* = 3.33). However, post hoc tests did not yield significant pairwise differences (all *p_Holm_* ≥ 0.086). For RNT responders (n = 21), the effect of Probe Direction was more pronounced (*F*(3, 60) = 4.88, *p* = 0.004, *η^2^* = .196, *BF₁₀* = 9.11). Post hoc analyses reproduced the gradual decrease in ERP amplitude from +90° (*M* = 3.77, *SEM* = 0.29) to −45° (*M* = 3.32, *SEM* = 0.30, *p_Holm_* = .028) observed in the full sample (Figure 5). As before, the −90° condition (*M* = 3.77, *SEM* = 0.29) deviated from this graded pattern, showing larger amplitudes than −45° (*p_Holm_* = .002).

### 4.3 The influence of NF training on alpha lateralization in the auditory attention task

If neurofeedback training has a lasting effect on task-based alpha lateralization, we would expect alpha power in the spatial pitch discrimination task after neurofeedback to be biased in the direction of the training condition (LNT; RNT). To test this prediction, we analyzed alpha power lateralization in the cue-to-target interval (0–1.9 s after cue onset) of the pitch discrimination task (see Figure 6).

**Figure 6.**
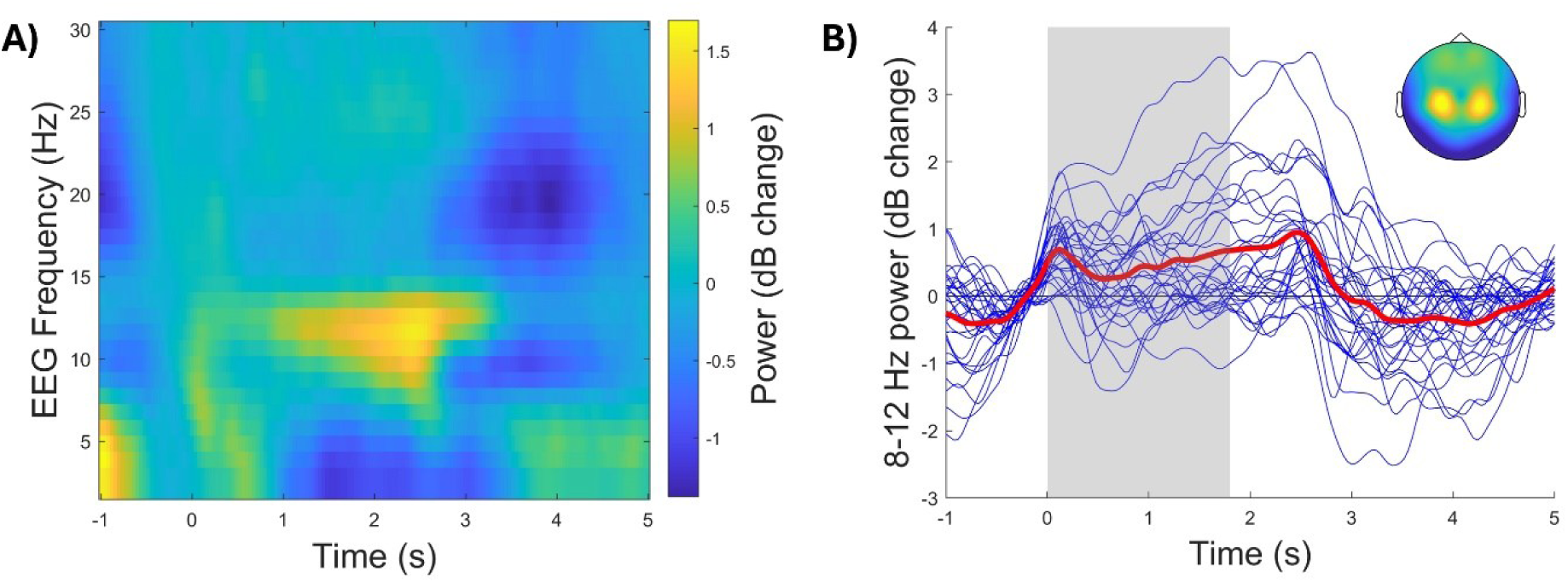
(A) Time-frequency representation of EEG power changes (in dB) relative to baseline (-0.5 to 0 s), averaged across all participants and electrodes, showing an increase in power in the 8–12 Hz alpha band following cue onset. (B) Individual (blue) and average (red) time courses of 8–12 Hz power changes over time, with a shaded region indicating the analysis window. The inset scalp map displays the topographical distribution of alpha power changes during the highlighted time window.

To assess the effects of neurofeedback during lateral target selection and lateral distractor suppression, we computed the Alpha Modulation Index (AMI) separately for conditions involving lateral target selection and lateral distractor suppression. Subsequently, the corresponding Alpha Asymmetry Index (AAI) was derived by subtracting the AMI values of the left and right electrode sets (see Methods).

First, we evaluated task-based alpha modulation during lateral target selection and lateral distractor suppression prior to neurofeedback. In line with previous findings (Wöstmann et al., 2019), we found a significant interaction between Task Condition (selection; suppression) x Hemisphere (left; right) (*F(*1, 33) = 12.60, *p* = 0.001, *η²ₚ* = 0.276, *BF_10_* = 447.26), indicating that target-selection-independent alpha power modulation contributes to distractor suppression. Post-hoc tests revealed larger AMI values over the left (*M*=0.019 *SEM*=0.009) than the right hemisphere (*M*=0.012 *SEM*=0.008) during lateral target selection (*p_holm_* = .003). During lateral distractor suppression, AMI did not differ between hemispheres (left: *M* = −0.013, *SEM* = 0.007; right: *M* = −0.004, *SEM* = 0.008; p*_holm_* = .364).

To assess the impact of neurofeedback on alpha power lateralization during target selection, we conducted a repeated-measures ANOVA with Time (pre-training; post-training) and Training (LNT; RNT) as within-subjects factors, and AAI_selection_ as the dependent variable. This analysis revealed no significant main effects of Time (*F*(1, 33) = 0.894, *p* = 0.351, *η²ₚ* = 0.026, *BF_10_* = 0.283) or Training (*F*(1, 33) = 0.067, *p* = 0.798, *η²ₚ* = 0.002, *BF_10_* = 0.231), and no significant Time × Training interaction (*F*(1, 33) =0.002, *p* = 0.962, *η²ₚ* < 0.001, *BF_10_* = 0.247). Together, these results indicate no evidence that neurofeedback systematically modulated tasked-based alpha lateralization during target selection (see Figure 7A – C).

**Figure 7.**
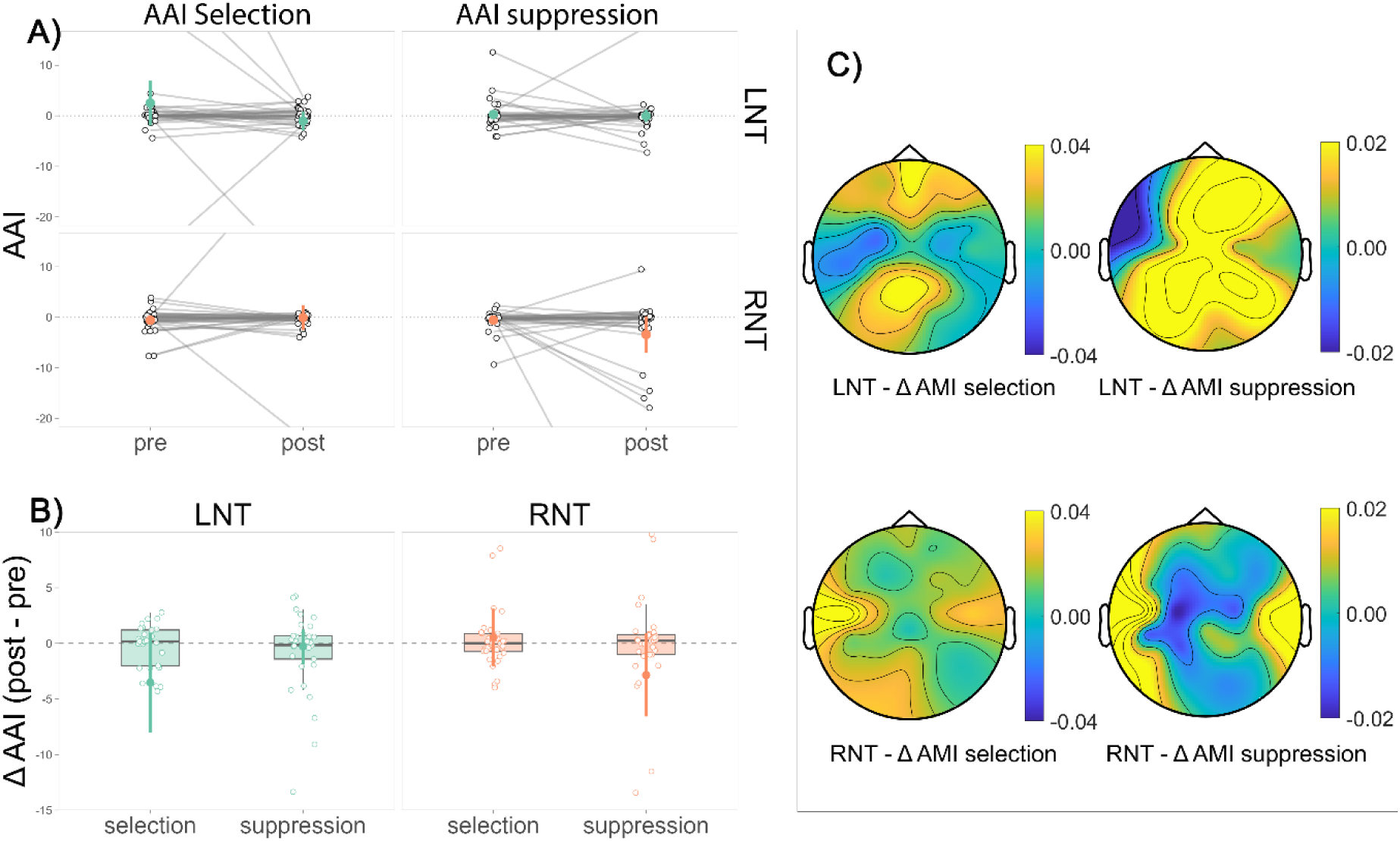
Effects of neurofeedback training on task-based alpha lateralization during the auditory attention task. (A) Alpha asymmetry index (AAI) during target selection (left column) and distractor suppression (right column) before (pre) and after (post) neurofeedback training, plotted separately for the LNT (green) and RNT (orange). Thin grey lines connect individual participants; thick colored markers show the group mean ± 95 % CI. (B) Change in AAI (post – pre) for the same conditions and groups (boxplots with individual data points and 95 % CI of the mean). (C) Scalp topographies of the change in alpha modulation index (ΔAMI, 8–12 Hz, 0–1.9 s) illustrating spatial distributions of training-related changes for target selection and distractor suppression in each group (LNT top row, RNT bottom row). Positive values reflect stronger alpha modulation following neurofeedback relative to the pre-neurofeedback baseline. Color scale represents dB change.

To examine whether neurofeedback changed alpha power lateralization during distractor suppression, we conducted a repeated-measures ANOVA with Time (pre-training; post-training) and Training (LNT; RNT) as within-subjects factors, using AAI_suppression_ as the dependent variable. This analysis revealed no significant main effects of Time (*F*(1, 33) = 0.176, *p* = 0.678, *η ²ₚ* = 0.005, *BF_10_* = 0.234) or Training (*F*(1, 33) = 0.032, *p* = 0.860, *η ²ₚ* = 0.001, *BF_10_* = 0.237), and no significant Time × Training interaction (*F*(1, 33) = 1.513, *p* = 0.227, *η ²ₚ* = 0.044, *BF_10_* = 0.635). These results suggest that NF training did not influence alpha lateralization during distractor suppression.

As before, we conducted a responder analysis of the task data because not all participants were able to modulate alpha lateralization during neurofeedback. To examine whether neurofeedback altered task-related alpha lateralization in training responders, we conducted exploratory repeated-measures ANOVAs separately for LNT and RNT responders, with Time (pre-training vs. post-training) as a within-subject factor. For selection, LNT responders showed no pre–post change (*F*(1, 21) = 2.94, *p* = 0.101, *η^2^_p_* = 0.123, *BF₁₀* = 1.26). RNT responders likewise showed no effect (*F*(1, 20) = 1.53, *p* = 0.230, *η^2^_p_* = 0.071, *BF₁₀* = 0.59). For suppression, LNT responders showed no reliable pre–post change (*F*(1, 21) = 1.21, *p* = 0.283, *η^2^_p_* = 0.055, *BF₁₀* = 0.55) and RNT responders showed no effect either (*F*(1, 20) = 0.06, *p* = 0.814, *η^2^_p_* = 0.003, *BF₁₀* = 0.30). Taken together, the responder analyses provide no evidence that NF training changed alpha lateralization in the auditory attention task - either for selection or suppression - converging with the conclusions from the registered all-participants analyses.

### 4.4 Effects of neurofeedback on gaze direction

As shifts in alpha lateralization have been found to co-occur with shifts in eye gaze (Liu et al., 2022a, 2023), we examined whether neurofeedback training biases the distribution of horizontal microsaccades toward the hemifield in which alpha power was enhanced. We conducted two separate analyses. The first focused on eye movement data recorded during neurofeedback to assess whether Training (LNT; RNT) differentially affected the distribution of microsaccades toward the left and right visual hemifields. The second analysis compared microsaccade distributions before and after neurofeedback. These analyses were conducted separately because the average microsaccade rate cannot be directly compared between neurofeedback and resting-state EEG. This is due to the different visual displays: during neurofeedback, participants viewed a computer game with a moving spaceship, whereas in the resting-state blocks, they fixated on a stationary cross.

First, we conducted a repeated-measures ANOVA including the within-subject factors Training (LNT; RNT) and Visual Hemifield (left; right), and microsaccade rate during neurofeedback as dependent variable. Results showed a significant main effect of Visual Hemifield (*F*(1,33) = 5.71, *p* = 0.023, η*²_p_* = 0.147, *BF_10_* = 2.303), with a higher microsaccade rate in the right visual hemifield compared to the left. This rightward bias in microsaccade rate aligns with the observed right-hemispheric alpha lateralization across participants. Contrary to our expectations, there was no evidence that the Training condition induces a gaze bias toward either visual hemifield during neurofeedback. Analyses revealed neither a main effect of Training (*F*(1,33) = 0.62, *p* = 0.437, *η^2^_p_* = 0.018, *BF_10_* = 0.41), nor a Training x Visual Hemifield (*F*(1,33) = 0.25, *p* = 0.617, *η^2^_p_* = 0.008, *BF_10_* = 0.240) interaction. A responder analysis conducted separately for LNT and RNT responders revealed numerically similar, though slightly larger, effect sizes for the main effect of Visual Hemifield in LNT (*F*(1,22) = 2.69, *p* = 0.115, *η^2^_p_* = 0.109, *BF_10_* = 0.81) compared to RNT responders (*F*(1,20) = 1.83, *p* = 0.192, *η^2^_p_* = 0.084, *BF_10_* = 0.59). This is somewhat surprising, as one would expect the opposite, namely that changes in alpha lateralization go along with increased microsaccade rates in the ipsilateral visual hemifield.

To test whether neurofeedback altered the distribution of horizontal microsaccades from before to after training, we conducted a repeated-measures ANOVA with the within-subject factors Time (pre-training rest; post-training rest), Training (LNT; RNT), and Visual Hemifield (left; right), using microsaccade rate as the dependent variable. Consistent with the earlier analysis, we found a significant main effect of Visual Hemifield (*F*(1,33) = 27.61, *p* < 0.001, *η^2^_p_* = 0.456, *BF_10_* = 2243), reflecting a higher microsaccade rate toward the right hemifield (*M* = 0.773, *SEM* = 0.051) compared to the left hemifield (*M* = 0.663, *SEM* = 0.042). Importantly, we observed a significant Training x Visual Hemifield interaction (*F*(1,33) = 4.36, *p* = 0.045, *η^2^_p_* = 0.117, *BF_10_* = 1.326) and a statistical trend for a three-way Training x Visual Hemifield x Time interaction (*F*(1,33) = 3.80, *p* = 0.060, *η ^2^* = 0.103, *BF*_10_ = 1.496) (see Figure 8). However, post hoc comparisons revealed no significant differences between Training conditions within either hemifield (left: *p_holm_* = .626; right: *p_holm_* = .176). In addition, a significant Time x Visual Hemifield interaction (*F*(1,33) = 4.63, *p* = .039, *η²ₚ* = 0.123, *BF_10_* = 1.784) indicated that microsaccade direction changed from before to after neurofeedback. Posthoc tests showed a pre–post difference in the right hemifield (*p_holm_* = .017), but not in the left hemifield (*p_holm_* = .836). Importantly, Bayesian evidence for the interaction effects was inconclusive (all *BF_10_* ≈ 1–1.8). Therefore, we conducted an additional responder analysis to examine whether these effects were driven by differential response patterns in LNT and RNT responders. This analysis revealed a significant Time x Visual Hemifield interaction in LNT responders (*F*(1,22) = 12.42, *p* = .002, *η²ₚ* = 0.361, *BF_10_* = 14.41), but not in RNT responders (*F*(1,20) = 0.01, *p* = .909, *η²ₚ* = 6.67*10^-4^, *BF_10_* = 0.340). In LNT responders, this interaction was driven by a reduction of the microsaccade rate in the right Visual Hemifield from pre to post neurofeedback (*M_pre_* = 0.919; *SD_pre_* = 0.342; *M_post_*=0.713; *SD_post_* = 0.296; *t*(22) = 2.65, *p_holm_* = 0.059). Together, these findings suggest that there are indeed differences in gaze behavior between RNT and LNT responders pre to post neurofeedback.

**Figure 8.**
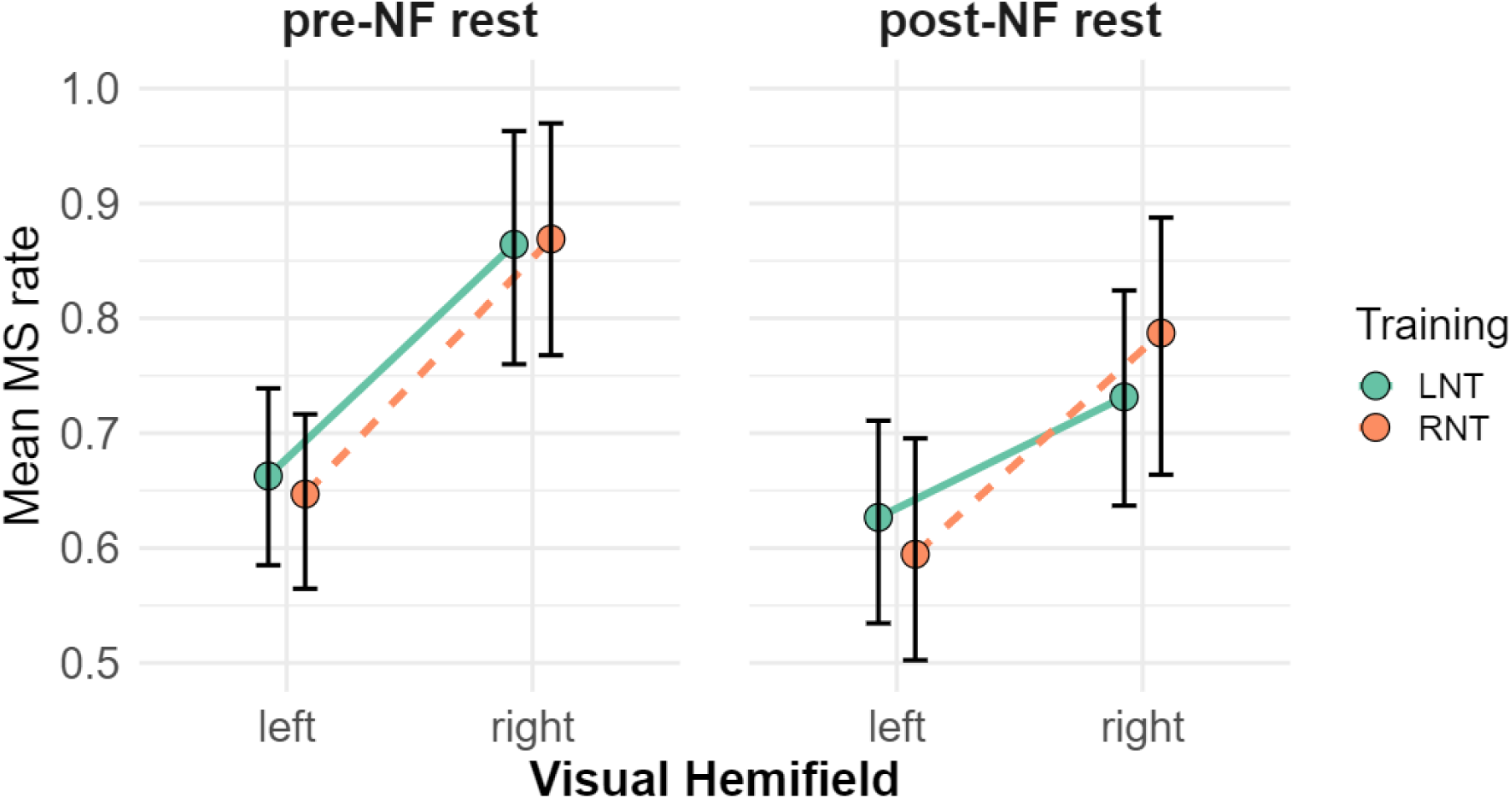
The influence of Training condition (LNT; RNT) on the distribution of horizontal microsaccades before and after neurofeedback. Microsaccade rate is plotted as a function of Visual hemifield, Time (pre-training rest, post-training rest), and Training condition (LNT; RNT). The figure shows a significant main effect of Visual Hemifield (*p* < .001, *BF_10_* = 2243), with a higher microsaccade rate toward the right compared to the left hemifield. Significant Training × Visual Hemifield (*p* = .045, *BF_10_* = 1.326) and Time × Visual Hemifield (*p* = .039, *BF_10_* = 1.784) interactions indicate that the training condition differentially influenced the direction of microsaccades and that this direction changed from before to after neurofeedback.

## 5 Discussion

Using EEG-based neurofeedback, our pre-registered analyses revealed no evidence for the hypothesis that parieto-occipital alpha lateralization serves a cross-modal functional role in auditory spatial attention. Although participants successfully modulated their alpha lateralization in the trained direction — confirming voluntary control over this neural signal in a within-subject cross-over design — this online modulation did not translate into online or offline changes in neural measures of auditory spatial attention. These findings challenge the idea that alpha power lateralization plays a functional role in auditory selective attention. In contrast, we found that LNT and RNT differentially altered gaze behavior relative to the pre-measurement baseline. The selective effect on gaze behavior in training responders points to a dissociation between oculomotor and auditory attentional systems and raises important questions about the functional specificity of alpha-based neurofeedback interventions.

### 5.1 Neurofeedback training enabled online control of alpha lateralization

Our results show that neurofeedback training induced changes in alpha lateralization in the trained direction. During LNT, AAI values increased, indicating a shift in lateralization towards the left hemisphere, while during RNT, AAI values decreased, reflecting a shift towards the right hemisphere. These findings align with previous research showing successful online modulation of alpha lateralization (Bagherzadeh et al., 2020; Mennella et al., 2017; Okazaki et al., 2015).

However, our experimental approach differs from previous studies in important ways. First, we employed a within-subject crossover design in which the same participants completed both LNT and RNT on two separate days. This allowed us to examine whether the same individuals are capable of flexibly modulating their alpha lateralization in both directions, providing a stronger test of voluntary control than is possible in a between-subject design. Most previous studies have relied on between-subject designs (Bagherzadeh et al., 2020; Choi et al., 2010; Mennella et al., 2017; Okazaki et al., 2015; Quaedflieg et al., 2016), with the exception of Schneider et al. (2020).

Second, it is important to note differences in the task and instructions used during neurofeedback training. In our study, participants performed a task that did not require spatial attention. They were asked to control a spaceship through mental effort alone, without any explicit spatial attentional demands. In contrast, some studies explicitly instructed participants to covertly direct their attention towards the respective visual hemifield (Okazaki et al., 2015; Schneider et al., 2020), while in the majority of studies participants were unaware that the trained biomarker, alpha lateralization, is associated with spatial attention (Bagherzadeh et al., 2020; Choi et al., 2010; Mennella et al., 2017; Quaedflieg et al., 2016; Rosenfeld, 2000; Rosenfeld et al., 1995). However, only few studies have controlled, as we did, for the possibility that participants modulated alpha lateralization during neurofeedback through spatial attentional strategies such as shifts in gaze direction (Bagherzadeh et al., 2020; Okazaki et al., 2015; Schneider et al., 2020). Importantly, while these studies demonstrated that central fixation can be maintained, our results go a step further by showing that the observed alpha lateralization effects cannot be attributed to differences in visual exploration strategies between training conditions, as evidenced by the absence of significant differences in the distribution of microsaccades between the LNT and RNT conditions.

Approaches to determining the baseline alpha lateralization also varied across studies. While we, in line with several previous studies, derived the individual baseline from a resting-state (eyes open) recording obtained immediately prior to training (Bagherzadeh et al., 2020; Mennella et al., 2017; Quaedflieg et al., 2016; Ros et al., 2017), other studies determined the baseline during a task block (Okazaki et al., 2015; Schneider et al., 2020). Furthermore, whereas some studies, including the present one, assessed the baseline on the same day immediately prior to neurofeedback (Bagherzadeh et al., 2020; Mennella et al., 2017; Quaedflieg et al., 2016; Ros et al., 2017), others obtained baseline measurements in a separate session on a preceding day (Rosenfeld et al., 1995; Schneider et al., 2020).

### 5.2 Neurofeedback does not modulate online auditory processing

In our pre-registered analysis, we found no evidence that the neurofeedback training condition influenced auditory evoked responses to lateralized probe tones, suggesting that online modulation of alpha lateralization does not translate into differential processing of sounds in the trained versus untrained hemifield. This finding stands in contrast to prior evidence from the visual domain, where probe-evoked responses were suppressed in the hemisphere with upregulated alpha power following neurofeedback training (Bagherzadeh et al., 2020), repetitive TMS (Romei et al., 2010; Thut et al., 2011), and tACS (Radecke et al., 2023).

Two interpretations of this null result are conceivable. The first and stronger interpretation is that parieto-occipital alpha lateralization does not play a functional role in modulating auditory sensitivity. This would suggest that the gating by inhibition model (Jensen & Mazaheri, 2010), which predicts that increased alpha power leads to reduced cortical excitability and attenuated evoked responses in the ipsilateral hemisphere, may apply to the visual but not to the auditory domain (see also section 5.6 for an integrative discussion). A second, more cautious interpretation is that the effect of alpha power lateralization on auditory sensitivity is weaker in the auditory domain than in the visual domain, and that the degree of alpha lateralization achieved in the current study was not sufficient to produce a measurable effect. Although LNT successfully shifted alpha lateralization towards the left hemisphere, the overall lateralization index remained negative. It is therefore possible that the induced leftward shift was not sufficient to modulate online auditory processing. This interpretation is to some extent supported by the responder analysis, which revealed larger ERP differences between probe directions in RNT compared to LNT responders.

### 5.3 Neurofeedback training causes no offline effects in resting-state and attention-based alpha lateralization

Our results show that while neurofeedback training enabled successful online control of alpha lateralization, these changes did not produce stable offline effects at rest and during the spatial auditory attention task. This suggests that a single session of neurofeedback may be insufficient to induce persistent changes in alpha lateralization that have been reported for multi-session protocols (Choi et al., 2010; Mennella et al., 2017). The observation that alpha lateralization returned to pre-training levels in the post-training resting state is consistent with research indicating the transient nature of single-session neurofeedback effects (Quaedflieg et al., 2016; Schneider et al., 2020). For example, Quaedflieg et al. (2016) found that although alpha asymmetry was modified during training, these changes did not persist at one-week or one-month follow-up assessments. Similarly, Schneider et al. (2020), despite observing behavioral improvements, found no corresponding pre-post changes in alpha lateralization measured on the same day. In the literature, Hebbian plasticity and homeostatic plasticity have been discussed as candidate mechanisms underlying neurofeedback-induced changes in brain activity (Ros et al., 2014). Hebbian plasticity predicts that repeated co-activation of neurons during training strengthens synaptic connections and promotes persistent changes, while homeostatic plasticity predicts a paradoxical compensatory response whereby the brain shifts in the opposite direction immediately after the intervention, so-called rebound effects. At the group level, our results are consistent with homeostatic plasticity, as alpha lateralization returned to baseline in both training conditions following the intervention. Interestingly, although the correlations between individual online and offline neurofeedback effects did not reach statistical significance, they showed divergent patterns across training conditions. While the association was positive in the LNT condition, it was negative in the RNT condition, suggesting that participants with the largest online training effects showed the smallest offline changes following RNT, which is indicative of a rebound effect.

### 5.4 Neurofeedback has no sustained effect on task-based alpha lateralization during auditory selective attention

Although our paradigm successfully elicited task-based alpha modulation and lateralization during the cue-to-target interval, we found no offline effect of neurofeedback training on these neural signatures, either in response to lateral targets or lateral distractors. This may not be entirely surprising given that neurofeedback-driven changes in alpha lateralization did not influence online auditory processing during training and did not persist in the offline resting-state EEG. This null finding is in line with Schneider et al. (2020), who similarly observed no pre-post changes in alpha lateralization despite significant behavioral improvements. Unlike Schneider et al. (2020), however, we were unable to evaluate behavioral task performance directly, as task difficulty was adaptively adjusted throughout the experiment to maintain balanced task demands. Our null result does stand in contrast to neurofeedback studies in the visual domain (Bagherzadeh et al., 2020; Okazaki et al., 2015), which reported sustained neural and behavioral biases in visual attention outlasting the training period. Indeed, the functional role of parieto-occipital alpha oscillations may differ between the visual and auditory domains (see also section 5.6). Recent large-scale work challenges the assumption that alpha lateralization drives auditory spatial filtering. Tune et al. (2021), in a study of 155 participants, demonstrated that while selective attention robustly modulated parieto-occipital alpha lateralization, these fluctuations did not directly predict trial-by-trial listening performance. Consistent with this, Getzmann et al. (2020) observed that attentional-shift-related increases in alpha lateralization were not directly coupled to behavioral accuracy. Taken together, these findings indicate that in the auditory domain, alpha lateralization may index attentional orienting without reliably translating into performance benefits.

### 5.5 Neurofeedback training affects gaze behavior differently in LNT and RNT responders

During neurofeedback training, gaze behavior did not differ between LNT and RNT conditions. Offline, however, LNT and RNT differentially affected resting-state gaze behavior in training responders, with LNT reducing the pre-existing rightward gaze bias from pre- to post-measurement.

The lack of online gaze differences indicates that participants did not adopt systematically different oculomotor strategies across training conditions, which is consistent with previous work (Bagherzadeh et al., 2020). This is relevant given recent work establishing a robust functional link between microsaccades and alpha lateralization. Specifically, Liu et al. (Liu et al., 2022b, 2023) demonstrated that spontaneous microsaccades are sufficient to drive transient lateralization of EEG alpha power, and Popov et al. (2022) showed that alpha power lateralization during auditory spatial attention is closely coupled with oculomotor activity and topographic gaze orientation, suggesting that spatial attention engages the oculomotor system in a manner analogous to the retinotopic organization of visual attention. Importantly, it has been argued that participants in neurofeedback studies might exploit covert spatial attention strategies, which bias microsaccade direction, to manipulate alpha lateralization rather than alpha modulation being the primary driver of attentional shifts (Gundlach & Forschack, 2020). Contrary to this concern, our data provide no indication that participants employed such strategies to alter their alpha lateralization. If anything, LNT responders showed a stronger gaze bias toward the right visual hemifield than RNT responders, a finding that is somewhat surprising. Given that LNT shifted alpha lateralization toward the left hemisphere, one would theoretically expect this to be accompanied by an increased microsaccade rate toward the left visual hemifield rather than the right.

Importantly, our results indicate that neurofeedback training induces offline changes in gaze behavior that differ as a function of training direction. Specifically, LNT reduced the gaze bias toward the right visual hemifield from pre- to post-neurofeedback during the resting state. This finding is consistent with previous evidence showing that neurofeedback training of alpha lateralization induces a persistent fixation bias in a subsequent free-viewing task in the direction of training (Bagherzadeh et al., 2020). Our observation of a pre-to-post shift in resting-state gaze bias suggests that, while the training did not produce sustained neural changes in alpha lateralization, it may have subtly re-tuned the homeostatic set-point of the oculomotor system, particularly in those who responded to the leftward training direction.

### 5.6 Is parieto-occipital alpha lateralization really a crossmodal switch for spatial attention?

In this pre-registered study, we found that volitionally changing neural alpha lateralization did not modulate online or offline neural signatures of auditory spatial attention. In contrast, LNT and RNT differentially affected gaze behavior in neurofeedback responders, which is consistent with previous research demonstrating online and offline effects of neurofeedback-induced changes in alpha lateralization on visual spatial attention (Bagherzadeh et al., 2020; Okazaki et al., 2015), as well as a link between alpha lateralization and gaze behavior (Liu et al., 2022b, 2023). The selective effect on gaze behavior in training responders points to a dissociation between oculomotor and auditory attentional systems, and raises important questions about the functional specificity of alpha-based neurofeedback interventions.

The prevailing theory holds that alpha-band lateralization (8-12 Hz) acts as a causal gate across sensory modalities by actively suppressing processing at unattended locations, thereby facilitating selective auditory attention (Frey et al., 2015; Jensen & Mazaheri, 2010). Brain stimulation studies have provided behavioral evidence supporting the role of parieto-occipital alpha lateralization in auditory spatial filtering (Deng et al., 2019; Wöstmann et al., 2018). However, the present findings speak against this account. If parieto-occipital alpha lateralization would provide truly a crossmodal attentional gate, its volitional modulation would be expected to directly influence the neural and electrophysiological correlates of auditory attention. Yet the neurofeedback-induced shift in alpha lateralization neither biased auditory processing nor produced lasting offline changes in alpha lateralization during auditory spatial attention. These findings are consistent with recent work reporting no reliable association between parieto-occipital alpha lateralization and auditory attention performance (Getzmann et al., 2020; Klatt et al., 2020; Tune et al., 2021).

A key limitation of the present study concerns the spatial specificity of the neurofeedback training. Because training was conducted in sensor space rather than source space, the precise cortical regions targeted cannot be determined with certainty. Nevertheless, given that parieto-occipital sensors were used, the intervention most likely modulated visual and parietal alpha activity rather than the auditory tau rhythm, which originates in the temporal lobe and is considerably weaker in amplitude and more difficult to detect at the scalp (Lehtelä et al., 1997; Wisniewski et al., 2024).

In summary, the present results challenge the notion that parieto-occipital alpha lateralization serves as a universal crossmodal spatial switch, and instead point to a functional dissociation between visual and auditory attention networks. Similar to the dissociation between spatial and temporal attentional networks (Coull & Nobre, 1998; Nobre & van Ede, 2018; Preisig & Meyer, 2025).

### 5.7 Conclusion

In conclusion, the present study demonstrates that neurofeedback training enabled online control of alpha lateralization, yet this did not translate into modulation of online auditory processing. Furthermore, neurofeedback training produced no offline effects on resting-state or attention-based alpha lateralization, nor did it exert sustained effects on task-based alpha lateralization during auditory selective attention. Taken together, these findings suggest that while participants were able to volitionally regulate their alpha lateralization during training, this did not generalize to broader neural or perceptual changes. A notable exception, however, concerns gaze behavior: neurofeedback training induced differential pre-to-post changes in microsaccade distribution in LNT and RNT responders, suggesting that the two training directions may have subtly re-tuned the oculomotor system in distinct ways, even in the absence of sustained effects on alpha lateralization itself.

## 6 Data and Code Availability

The raw data and the final analysis code supporting the findings of this study will be made publicly available upon acceptance for publication on osf.io.

## 7 Author Contributions

F.S.: methodological implementation, data acquisition, data analysis, writing - original draft, writing - review & editing.

T.R.: conceptualization, validation, writing - review & editing.

B.C.P.: conceptualization, methodological implementation, validation, data analysis, writing - original draft, writing - review & editing, funding acquisition.

## 10 Supplementary Material

No supplemental material

## 11 Acknowledgements

We thank Anouk Glättli and Nataliya Fartdinova for their assistance, and Andrew Clark, Omid Marxen, and Bernhard Wandernoth for technical assistance and software development.

